# Cytochrome *b*_6_*f* complex inhibition by antimycin-A requires Stt7 kinase activation but not PGR5

**DOI:** 10.1101/2020.08.22.262592

**Authors:** Felix Buchert, Michael Hippler

## Abstract

Ferredoxin-plastoquinone reductase (FQR) activity during cyclic electron flow (CEF) was first ascribed to the cytochrome *b*_6_*f* complex (*b*_6_*f*). However, this was later dismissed since *b*_6_*f* inhibition by antimycin-A (AA) could not be reproduced. AA presumably fails to ligate with haem *b*_h_, at variance with cytochrome *bc*_1_ complex, owing to a specific Qi-site occupation in *b*_6_*f*. Currently, PROTON GRADIENT REGULATION5 (PGR5) and the associated PGR5-Like1 are considered as FQR in the AA-sensitive CEF pathway. Here, we show that the *b*_6_*f* is conditionally inhibited by AA in a PGR5-independent manner when CEF is promoted. AA inhibition, demonstrated by single *b*_6_*f* turnover and electron transfer measurements, coincided with an altered Qi-site function which required Stt7 kinase activation by a strongly reduced plastoquinone pool. Thus, PGR5 and Stt7 were necessary for *b*_6_*f* activity and AA-sensitive electron transfer in CEF-favouring conditions. Extending previous findings, a new FQR activity model of the *b*_6_*f* is discussed.

## Introduction

Light is captured by two photosystems (PSI and PSII) and their associated light harvesting complexes (LHCI and LHCII) which results in the splitting of water by PSII and the reduction of ferredoxin (Fd) by PSI. Reduced Fd carries the electrons to Fd-NADP(H) oxidoreductase (FNR) which generates NADPH in this linear electron flow (LEF) process, thus providing the reducing equivalents for CO_2_ fixation in the Calvin Benson cycle. Light-driven charge separation and water splitting generate a membrane potential (ΔΨ) and a proton gradient (ΔpH), respectively, and this proton motive force (pmf) drives ATP synthesis. Both photosystems are functionally connected by the cytochrome *b*_6_*f* complex (*b*_6_*f*) which sophisticatedly transfers electrons from plastoquinol (PQH_2_) to plastocyanin^1, 2^. These steps involve an electron bifurcation within the stromal PQH_2_ binding pocket, the Qo-site. The first donated electron from PQH_2_ enters the high-potential chain and the second one enters the low-potential chain. The proton release/binding that is associated with the interconversion of PQH_2_ to plastoquinone (PQ) generates a ΔpH, and charge separation within the low-potential chain produces ΔΨ. Besides minor subunits (PetG, L, M and N), the *b*_6_*f* core subunits are cytochrome *f* (cyt.*f*), the Rieske iron sulphur protein (ISP), subunit-IV and cytochrome *b*_6_. According to their redox midpoint potential (E_m_), which is between +300 mV (Rieske ISP) and +380 mV (cyt.*f*), the high-potential chain differs from the low-potential chain. The latter is formed by three redox cofactors: haems *b*_l_ (E_m_ = −130 mV), *b*_h_ (E_m_ = −35 mV), and *c*_i_. Haem *c*_i_ is near haem *b*_h_ in the stromal Qi-site and is linked to cytochrome *b*_6_ via a single thioether bond, lacking amino acid axial ligands. Its E_m_ ranges from +100 mV to approximately −150 mV, when ligated to semiquinone analogues^3^. Moreover^4^, the presence of a ΔΨ may modulate the E_m_ of haem *c*_i_, resulting in the shared electron to reside as *b*_h_^red^/*C*_i_^ox^. The structural properties of haem *c*_i_ were suggested by some authors^5, 6^ to be relevant for cyclic electron flow (CEF) where Fd reduces haem *c*_i_ and thereby equips the *b*_6_*f* with Fd-PQ reductase (FQR) activity. CEF between PSI and the *b*_6_*f* is important in ATP-depleted conditions to drive the Calvin Benson cycle by compensating for the excess of NADPH, and by protecting PSI from photodamage. The latter is realized by various processes such as photosynthetic control of the *b*_6_*f* that slows down PQH_2_ oxidation in the Qo-site at low lumen pH.

The light capturing efficiency may be unevenly distributed within the photosynthetic machinery since light quality is dynamically changing. Therefore, lateral movements of mobile LHCII transiently serve as compensatory adjustments between PSII and PSI, which is termed state transitions^7^. A crucial sensor/responder function for state transitions is the PQ pool redox state and a thylakoid-associated Ser/Thr protein kinase. In Chlamydomonas reinhardtii, the kinase is termed Stt7 (STN7 in Arabldopsls fhallana) and it is activated by a reduced PQ pool. Various phosphatases were identified as kinase antagonists and, in order to phosphorylate LHCII and favour its movement towards PSI, Stt7 activation requires a functional *b*_6_*f* Qo-site^8^. The kinase interacts with the *b*_6_*f*^9^ and it is not known whether Stt7 is required for the *b*_6_*f* function although it phosphorylates various residues in subunit-IV and in the loosely attached subunit PETO^10, 11, 12^. The latter is an algal CEF effector protein that was identified in a PSI-*b*_6_*f* supercomplex^13, 14, 15, 16^. FNR, which is considered as a *b*_6_*f* subunit^17, 18, 19^, was routinely detected in these enriched fractions together with proteins of the CEF pathway that depends on PROTON GRADIENT REGULATION5 (PGR5). We recently found that the algal *b*_6_*f* failed to operate only in CEF-promoting conditions when PGR5 was absent, which we attributed to a dysfunctional FQR activity of the *b*_6_*f*^20^. This implied that the *b*_6_*f* operates via a canonical Q cycle in LEF conditions and as FQR during CEF for which PGR5 ensures Fd supply by recruiting FNR to the membrane^20, 21^. Our findings challenged an acknowledged model that ascribes the FQR to be formed by PGR5/PGR5-like1^22, 23^. The PGR5 interactome was mainly discovered owing to the observation^24^ that the *b*_6_*f* was not inhibited by antimycin-A (AA), unlike PGR5-dependent slow chlorophyll fluorescence kinetics *in vitro*^22^. AA is a well-known Qi-site inhibitor in the respiratory cytochrome *bc1* complex^25^. Initial studies demonstrated inhibition of the *b*_6_*f* by AA^26^ which, at the time, supported the view that the *b*_6_*f* acts as FQR during Fd-dependent cyclic photophosphorylation^27, 28, 29^.

Here, we show *in vivo* that the *b*_6_*f* is exclusively inhibited by AA in CEF-favouring conditions as a function of the PQ pool redox state. The inhibition was independent of PGR5 but relied on Stt7 kinase activity. Efficient electron transfer under CEF-promoting conditions relied on the AA sensitive step of photosynthesis, i.e. on the FQR activity of the *b*_6_*f*. A refined model of our previous findings is discussed where Stt7 primes the Qi-site and PGR5 provides Fd.

## Results

### Conditional antimycin-A sensitivity of the cytochrome *b*_6_*f* complex

Based on the structure of the algal *b*_6_*f* with the unusual Qi-site occupation^5^ and the ligation of AA to haem-bh in the respiratory cytochrome *bc1* complex^25^, it has been proposed that AA fails to act as Qi-site inhibitor in *b*_6_*f*^2^. The proposal has been reinforced by various conflicting reports on the *b*_6_*f* inhibitor potency of AA in isolated chloroplasts from vascular plants^24, 27, 28, 29^. To test AA efficacy *in vivo*, we used the green alga *Chlamydomonas reinhardtii* grown at moderate light. Indeed, we did not observe an effect of AA on the *b*_6_*f* kinetics in light-adapted cells that were mixed thoroughly throughout the measurement, i.e. kept in an oxic state (Fig. 1a). In both control and AA sample, the majority of photo-oxidized cyt.*f* was re-reduced within 10-ms, and the *b*-haems net reduction and oxidation was finished ~2-ms and ~30-ms after the flash, respectively. The redox amplitudes of the *b*-haems signals were slightly increased in the presence of AA and the reactions ceased at pre-flash levels, like in the control. When monitoring the electrochromic shift (ECS) of the photosynthetic pigments in these samples, the electrogenic contribution of the *b*_6_*f* during the first 10-ms after the flash (*b*-phase) was hardly visible in the control of Fig. 1b. As expected in light-adapted cells, the *b*-phase was masked by the coinciding and very efficient ΔΨ consumption via ATP synthase (in the thioredoxin-mediated reduced state). The *b*-phase was more apparent when AA was present, owing to a slowdown of ATP synthesis. Again, this was expected since the inhibitory AA impact on respiration interfered with metabolic coupling between chloroplasts and mitochondria^30, 31^. Thus, the exchange of reducing equivalents from the plastid for mitochondrial ATP resulted in a more reduced, slightly ATP-depleted chloroplast which is known to slowdown algal ATP synthesis^32^. The addition of PSII inhibitors to oxic AA samples revealed a pronounced *b*-phase that coincided with a further slowdown of the ATP synthase activity (grey symbols in Fig. 1b), pointing to an even more ATP-depleted chloroplast. Next, we monitored electron transfer rates during a 10-s illumination period (Fig. 1c). Since the samples were light-adapted, the electron sink capacity of the Calvin Benson cycle was still active after 30-s darkness at the onset of re-illumination. Accordingly, initial rates of ~175 charge separations/s/PSI levelled off in the controls after ~500-ms of illumination which yielded quasi steady state rates of ~75 charge separations/s/PSI. A transient slowdown which was followed by a re-acceleration was observed between 10-ms and 500-ms of illumination. In AA samples after the 30-s dark period, the initial rates of ~200 charge separations/s/PSI were slightly higher. Compared to controls, elevated rates were observed up to 50-ms of illumination. It is of note that a slight, AA concentration-dependent stimulation of non-cyclic photophosphorylation has been observed in earlier reports as well^29^. Following the AA-stimulated phase up to 50-ms, a period of diminished electron transfer was observed and a virtual steady state of ~75 charge separations/s/PSI was established no later than 2.5-s of illumination, owing to a slower re-acceleration phase. Compared to AA samples, only slightly lower initial electron transfer transients up to 25-ms of re-illumination were measured when PSII was inhibited via HA/DCMU in the presence of AA. In these strongly ATP-depleted samples after 30-s darkness, the light-induced electron transfer ceased exponentially within ~200-ms and yielded steady state rates of ~5 charge separations/s/PSI after ~500-ms light.

**Fig. 1.**
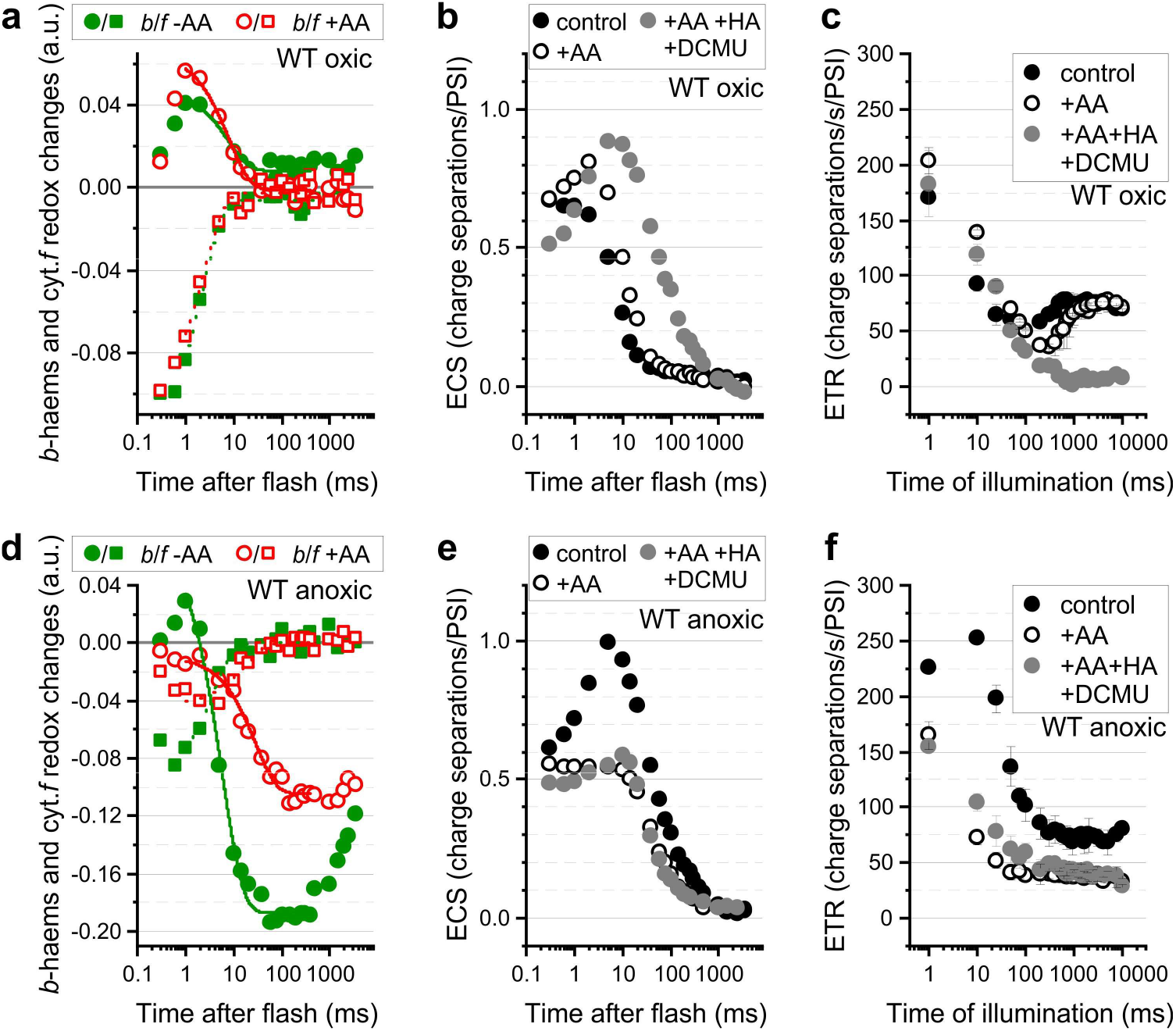
The conditional sensitivity of the cytochrome *b*_6_*f* complex to antimycin-A depends on a cellular redox poise. Here, light-adapted WT cells are shown that experienced 30-s darkness before the measurements. Representative kinetics and, where indicated, means of three biological replicates (± SD) are shown. **a** Redox signals for cytochrome *f* (cyt.*f*, squares) and *b*-haems (circles) are shown in oxic WT controls (-AA, closed symbols) and after treatment with antimycin (+AA, open symbols). The AA treatment did not substantially alter cyt.*f* re-reduction upon a flash, shown by a signal increase in the positive direction that finished no later than 50-ms. AA did also not affect the *b*-haems reduction (positive signals) and re-oxidation to pre-flash levels, finishing at ~100-ms. **b** The electrochromic shift (ECS) signals varied slightly since the *b*_6_*f*-dependent electrogenic 10-ms phase upon a laser flash was obscured in light-adapted controls (black) by a highly active ATP synthase that resulted in a fast ECS decay phase. Slowdown of the latter revealed the *b*_6_*f*-phase in AA samples (white), also upon PSII inhibition by addition of HA and DCMU (grey, see Material and Methods). **c** An ECS-based electron transfer rate (ETR) measurement during a 10-s re-illumination shows the gradual decrease of high initial ETRs to quasi steady state levels, established no later than 2.5-s. Unlike AA treatment, PSII inhibition lowered steady state ETR. See text for details. **d** Redox signals in the *b*_6_*f*, as in panel a, are shown for anoxic cells. While cyt.*f* in the controls was similar as in oxic samples, the *b*-haems oxidation phase reached signals below the pre-flash levels, followed a by a re-reduction at ~500-ms. AA treatment inhibited cyt.*f* reduction and *b*-haems redox kinetics. **e** ECS kinetics revealed the *b*_6_*f*-phase in the control due to slow ATP synthesis. AA treatment abolished the *b*_6_*f*-phase and light adaptation in the absence of PSII activity partially restored it (cf. white and grey 10-ms phases). **f** Re-illumination of anoxic controls sustained an elevated ETR up to 25-ms. The steady state ETR and the induction phase depended on AA-sensitive processes. The induction phase could be partially recovered when cells were illuminated in absence of PSII photochemistry.

A similar set of experiments is shown in Figs. 1d–1f for anoxic algae. In line with a previous report^20^, the *b*_6_*f* kinetics of the controls in Fig. 1d differed from oxic samples. cyt.*f* re-reduction was slightly delayed, finishing shortly after 10-ms darkness. The initial *b*-haems signals in anoxic controls showed a similar 1-ms net reduction after the flash. However, kinetics differed profoundly thereafter by displaying a large net oxidation phase up to 60-ms, followed by a *b*-haems re-reduction after ~500-ms darkness. In contrast to oxic samples, the presence of AA inhibited the *b*_6_*f* under anoxic conditions, both on the levels of cyt.*f* re-reduction and *b*-haems redox kinetics (cf. open symbols in Figs. 1a and 1d). The impaired *b*_6_*f* contribution to the light-dependent ΔΨ generation was also visible on the level of ECS kinetics after the flash, as shown by the absence of the *b*-phase upon AA treatment (cf. open circles in Figs. 1b and 1e). Since ATP synthase is slowed down in reducing conditions^32^, the *b*-phase became apparent in the controls of Fig. 1e. When AA-treated anoxic cells were further illumination upon PSII inhibition via HA/DCMU, the *b*-phase was partially restored (grey symbols in Fig. 1e). This suggests mitigation of the AA effect in absence of PSII photochemistry. We also checked electron transfer rates (Fig. 1f), clearly demonstrating the inhibitory effect of AA in anoxic cells. The controls displayed sustained initial transients with up to ~250 charge separations/s/PSI in the first 10-ms of re-illumination (at least). This highly active initial phase was followed by an exponential decrease in activity to reach the apparent steady state of ~75 charge separations/s/PSI after ~300-ms in the light. AA treatment diminished both the initial phase and the steady state. The rates in AA samples, starting from ~165 charge separations/s/PSI, levelled off at ~40 charge separations/s/PSI after 50-ms of light. Unexpectedly, slightly higher transients were observed between 10-ms and 200-ms of light when PSII was subsequently inhibited.

### Mitigated antimycin-A sensitivity of the cytochrome *b*_6_*f* complex in the *stt7-1* kinase mutant

The observations in Fig. 1 that show different AA effects in oxic and anoxic WT raised the question whether the PQ pool redox state governs the (partially reversible) AA sensitivity of the b6f, and whether active modulation via the Stt7 kinase is involved. To test this, we examined the *stt7-1* strain under anoxic conditions in Fig. 2 since the mutant is fully devoid of the kinase^10^. Interestingly in anoxic *stt7-1*, AA treatment had a milder inhibitory effect on *b*_6_*f* redox reactions after the flash (Fig. 2a). Accordingly, the *b*-phase in the ECS kinetics was less affected (Fig. 2b). When monitoring photosynthetic electron transfer during the 10-s illumination in the *stt7-1* control (Fig. 2c), the initial rate of ~240 charge separations/s/PSI resembled WT control (cf. Fig. 1f). The mutant, however, did not sustain high initial rates up to (at least) 10-ms light where rates have dropped to ~200 charge separations/s/PSI already. Thereby, *stt7-1* levelled off immediately to establish a quasi steady state of ~40 charge separations/s/PSI after ~200-ms in the light, which was lower than WT. The initial rates and the steady state in AA-treated *stt7-1* were very similar as in the control, with exception of slightly lower rates during the transition between 10-ms to 300-ms. To summarize the conditional AA effect in WT *b*_6_*f* and the data of *stt7-1*, the rate constants for cyt.*f* reduction (*k_f-red_*, Fig. 2d) and *b*-haems oxidation (*k_b-ox_*, Fig. 2e) are shown. This data served as an estimate as to whether the *b*_6_*f* was limiting during the induction phase of photosynthesis shown in Figs. 1c, 1f and 2c. Apart from the above-mentioned slight delay of re-reduction, *k*_*f*-red_ was not significantly altered in anoxic WT (~320 s^−1^) compared to oxic cells (~350 s^−1^). The presence of AA in oxic WT samples produced an insignificant accelerating effect on *k*_*f*-red_. Moreover, *k*_*f*-red_ in anoxic WT was faster than in the absence of Stt7 kinase (~190 s^−1^) and was severely slowed down in the presence of AA (~40 s^−1^). The AA-induced slowdown of *k*_*f*-red_ was insignificant in *stt7-1* (~150 s^−1^). As reported previously^20^, anoxic WT in Fig. 2e showed slightly faster *k*_*b*-ox_ (~160 s^−1^) compared to oxic samples (~100 s^−1^). Anoxic *stt7-1* (~90 s^−1^) was significantly slower than WT. AA did not produce a significant inhibitory effect on *k*_*b*-ox_ in *stt7-1* (~80 s^−1^) which stood in contrast with the AA inhibition in WT (~40 s^−1^).

**Fig. 2.**
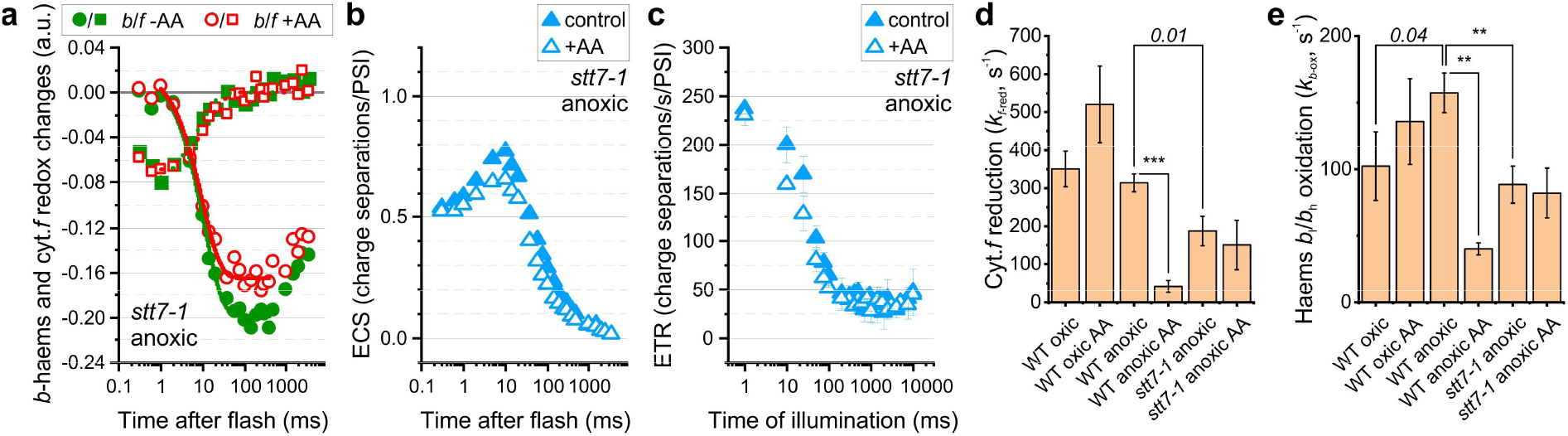
The antimycin-A sensitivity of the cytochrome *b*_6_*f* complex was less pronounced in anoxic *stt7-1* cells. The light-adapted samples were kept in darkness for 30-s before the measurements. Representative kinetics and, where indicated, means of three biological replicates (± SD) are shown. **a** The *bef* redox kinetics are shown and resemble anoxic WT signals of Fig. 1d. At variance with WT, AA treatment produced only a slight slowdown of cyt.*f* reduction and *b*-haems oxidation in *stt7-1*. **b** Accordingly, the electrogenic *b*_6_*f*-phase of the ECS signals was less affected by AA, **c** as well as the electron transfer rate (ETR) during re-illumination in the presence of AA. The *stt7-1* was less capable to sustain WT-like ETR after several ms of light and produced a lower steady state, compared to Fig. 1f. The *b*_6_*f* kinetics analysis of Figs. 1 and 2 is summarized in panels d and e (see Materials and Methods). Unless marked, *P* is indicated in italics (two-tailed Student’s t-test **P < 0.005 and ***P < 0.0005). **d** The cyt.*f* reduction rate constants (*k*_*f*-red_) showed that only anoxic WT was inhibited by AA and *k*_*f*-red_ in anoxic *stt7-1* controls was slower than WT. **e** The apparent oxidation rate constants for the *b*-haems (*k*_*b*-ox_) showed that anoxic condition had an accelerating effect in WT controls which coincided with AA susceptibility. In *stt7-1* anoxic controls, *k*_*b*-ox_ was slower than in WT and not significantly affected by AA.

### The cytochrome *b*_6_*f* complex in *pgr5* is sensitive to antimycin-A

After demonstrating AA inhibition in the anoxic WT *b*_6_*f*, and linking the effect to the Stt7 kinase and the PQ pool redox state, we followed our initial study^20^ and questioned whether the AA sensitivity of WT under CEF-favouring conditions is linked to PGR5 function. In other words, is the PGR5 polypeptide involved in forming the AA-sensitive FQR machinery, as suggested in a widely accepted model for this type of CEF^22, 23^? To test this, cells were grown in weak light to prevent PSI photodamage in pgr5^21, 33^. As expected, the low-light anoxic WT showed an AA effect on the level of *b*_6_*f* kinetics (Fig. 3a) as well as the electrogenic *b*-phase after the flash (Supplementary Fig. 1a). The same applied for electron transfer transients during photosynthetic induction (Fig. 3b), except that slightly higher apparent steady state rates of ~100 charge separations/s/PSI were obtained after ~300-ms in the WT control (cf. Fig. 1f for WT grown in moderate light). When measuring anoxic pgr5, we also observed a slowdown of *b*_6_*f* kinetics in the presence of AA (Fig. 3c). However, the *b*-haems oxidation amplitude was almost not diminished in pgr5 AA samples, unlike the electrogenic *b*-phase amplitude (Supplementary Fig. 1b). The control pgr5 failed to sustain efficient electron transfer rates which was evidenced here by the low virtual steady state of ~35 charge separations/s/PSI (Fig. 3d). Nevertheless, during the initial phase up to (at least) 10-ms of illumination, the mutant resembled WT and high rates of ~235 charge separations/s/PSI were observed in pgr5 controls. As expected from the AA effect on the *b*_6_*f* (Fig. 3c), electron flow was severely affected by the inhibitor in pgr5 (Fig. 3d), which resembled WT during the first 100-ms of light and levelled off at only ~25 charge separations/s/PSI.

**Fig. 3.**
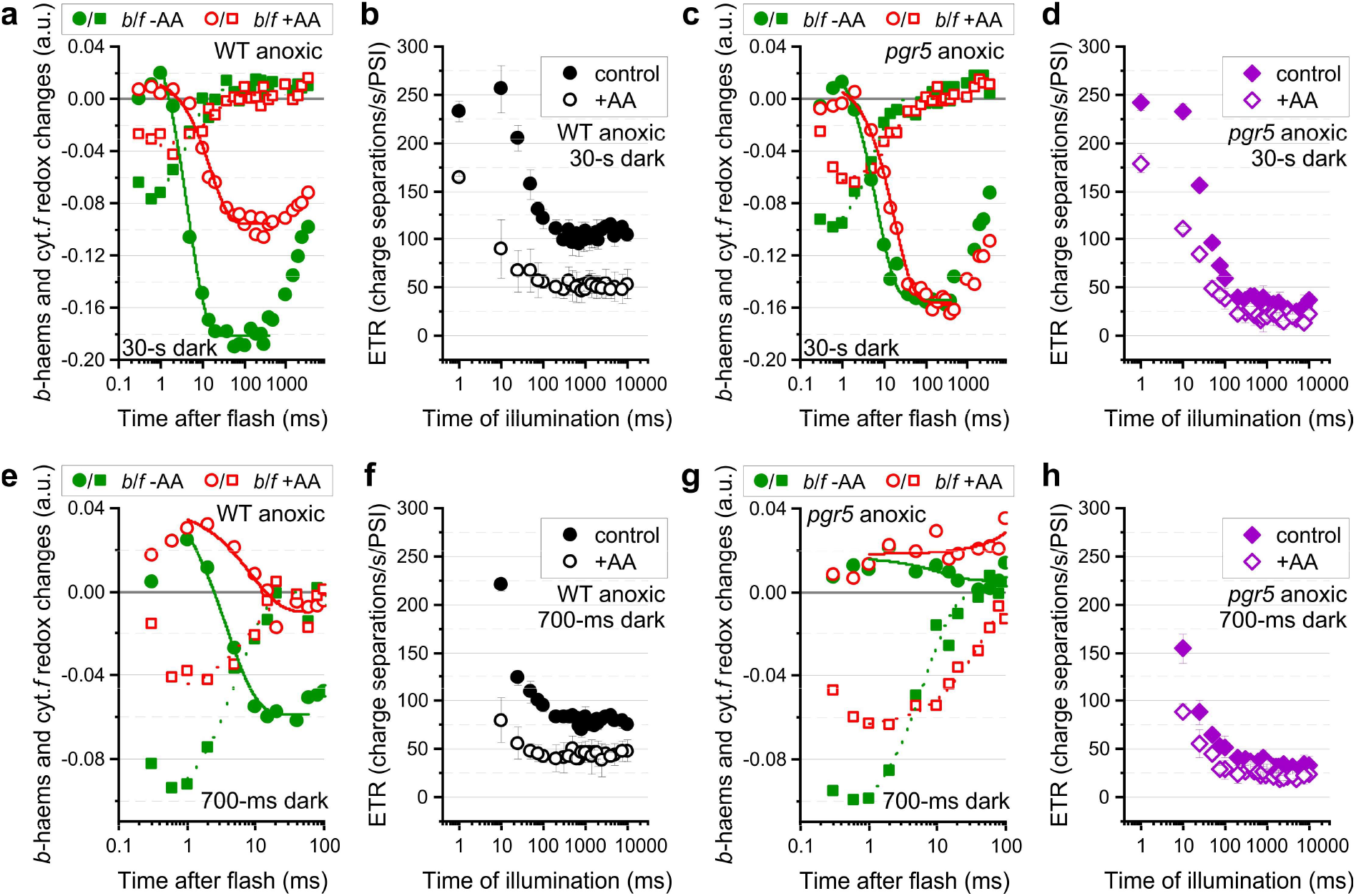
The conditional sensitivity of the cytochrome *b*_6_*f* complex to antimycin-A is displayed in *pgr5*. Light-adapted WT and mutant cells experienced both 30-s (panels **a–d**) and 700-ms darkness before the measurements (panels **e–h**). Representative kinetics and, where indicated, means of three biological replicates (± SD) are shown. **a** The *b*_6_*f* kinetics in anoxic WT and the AA effects resemble Fig. 1d despite different growth conditions. **b** This also applied for the light-induced electron transfer rate (ETR) although they developed a slightly higher steady state compared to Fig. 1f. **c** The *b*_6_*f* kinetics in anoxic *pgr5* resembled WT from panel a. Both, the cyt.*f* re-reduction and *b-* haems oxidation were slowed down by AA. At variance with WT, the *b*-haems amplitude was not altered in the presence of AA. **d** The mutant managed to produce WT-like ETR, at least during the first 10-ms of light, followed by a rapid ETR drop to a steady state lower than in WT. AA treatment had an effect on the induction phase, less on the steady state. **e** Compared to panel a, the short dark-relaxation of 700-ms resulted in slightly modified *b*_6_*f* kinetics in anoxic WT which mainly involved a smaller *b*-haems oxidation amplitude. **f** ETR after 10-ms light was almost as high as in panel b but it declined more rapidly with ongoing illumination. **g** The 700-ms *b*_6_*f* signals in *pgr5* differed from WT by showing slightly slower cyt.*f* reduction and almost no net redox changes of *b*-haems (for electrogenic events see Supplementary Fig. 1). **h** Compared to panel d, a significantly diminished ETR was seen in *pgr5* already with the first record at 10-ms light.

The 30-s dark period served to dark-relax the system before measurements, i.e. restore the availability of electron acceptors/donors, diminish the ΔpH-dependent electron flow downregulation, etc. However, various dark processes influence the PQ pool redox state such as the type II NAD(P)H dehydrogenase which has an activity of ~2 electrons/s^34, 35^. In the absence of oxygen, i.e. the substrate for plastid terminal oxidase, this should favour PQH_2_ production in the dark which may have influenced our measured initial electron transfer rates. To exclude this potential source of additional electrons in the chain before the measurements, the *b*_6_*f* and photosynthetic induction were analysed in the same WT and pgr5 samples after a short dark period of 700-ms as well. Compared to the 30-s dark ECS (i.e. ΔΨ), the values at 700-ms were similar in the controls (Supplementary Fig. 1c). After 700-ms, ΔpH was likely not completely collapsed. Likewise, re-equilibration of the redox carriers was still ongoing. Therefore, only the first 100-ms after the flash are shown for anoxic WT in Fig. 3e. The cyt.*f* re-reduction to pre-flash levels was finished no later than 20-ms during which the *b*-haems displayed a net oxidation, despite electrogenic charge transfer from haem *b*_l_^red^ to haem *b*_h_^ox^ (see inset in Supplementary Fig. 1c). AA treatment slowed down both kinetics (Fig. 3e). Re-illumination after the short dark period yielded high initial rates of ~225 charge separations/s/PSI in WT controls after 10-ms light, which levelled off at ~85 charge separations/s/PSI (Fig. 3f). The AA samples were identical, compared to 30-s dark (cf. Fig. 3b). After 700-ms darkness, the flash-induced *b*_6_*f* turnover in pgr5 controls differed from WT by showing a slightly slower cyt.*f* re-reduction and absence of large net redox changes in the *b*-haems (Fig. 3g) when electrogenic events took place in the *b*_6_*f* (see inset in Supplementary Fig. 1c). As shown in Fig. 3h, the initial rates after 10-ms re-illumination were not as high when pgr5 controls experienced the short dark period, yielding ~155 charge separations/s/PSI (cf. Fig. 3d). Again, AA treatment in pgr5 inhibited the induction and maintenance of photosynthetic electron transfer during the 10-s re-illumination.

To summarize the *b*_6_*f* characterization of low-light cultures in Fig. 3, Fig. 4a shows that the AA-induced slowdown of *k*_*f*-red_ after 30-s dark was similar in WT and pgr5. The corresponding *k*_*b*-ox_ are shown in Fig. 4b. In line with a previous report^20^, *k*_*b*-ox_ in the pgr5 control was slower than in WT (Fig. 4b). However, AA treatment yielded similar low *k*_*b*-ox_ in both strains. When cells experienced only 700-ms darkness in Fig. 4c, *k*_*f*-red_ was similar in WT controls (cf. Fig. 4a for 30-s dark). This was unexpected and suggests weak photosynthetic control in low-light WT cultures, which could explain the above-mentioned high initial rates of ~225 charge separations/s/PSI after 700-ms dark (Fig. 3f). However, *k*_*f*-red_ in pgr5 controls was slower than in WT after 700-ms darkness (Fig. 4c). The pgr5 *k*_*b*-ox_ parameter could not be obtained after 700-ms darkness (cf. Fig. 3g) since *b*-haems reduction and oxidation kinetics (see *b*-phase in Supplementary Fig. 1c) nullified net redox changes independently of AA.

**Fig. 4.**
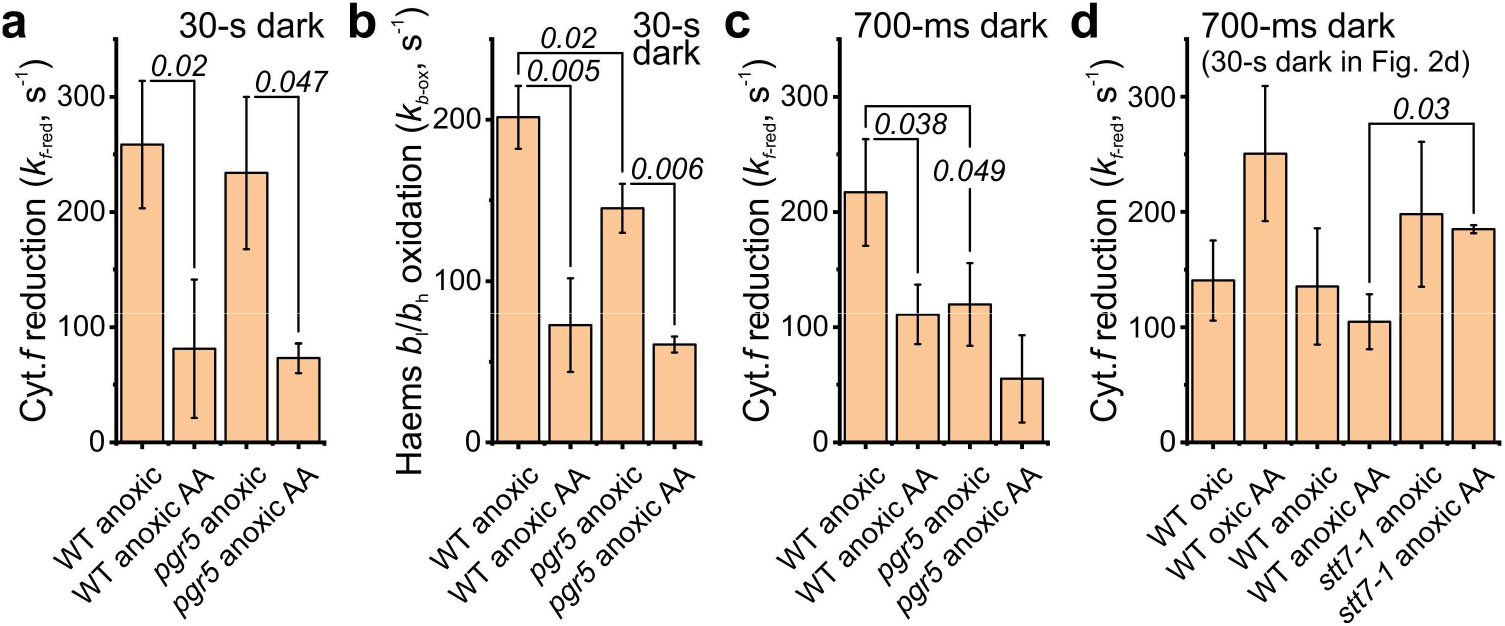
The *b*_6_*f* kinetics analysis of Fig. 3 is summarized and cyt.*f* reduction from samples in Figs. 1 and 2 are characterized after 700-ms darkness. Means of three biological replicates are shown (± SD) and *P* is indicated in italics (two-tailed Student’s t-test). **a** The cyt.*f* reduction rate constants (*k*_*f*-red_, see Materials and Methods) show that both strains from Fig. 3 were inhibited by AA after 30-s darkness to similar extents. **b** Accordingly, the apparent oxidation rate constants for the *b*-haems (*k*_*b*-ox_) showed the same trend. In pgr5 controls, *k*_*b*-ox_ was slower than in WT. **c** The *k*_*f*-red_ after 700-ms darkness were slower in pgr5 controls. A significant AA effect was seen in WT only. The *k*_*f*-red_ in WT controls was comparable to panel a. **d** The 700-ms *k*_*f*-red_ parameters of moderate light cultures are shown. The short dark period slowed down *k*_*f*-red_ in all controls, except the stt7-1 which was constantly slower unless AA was present (cf. Fig. 2d).

Independent of PGR5 but related to the above-mentioned weak photosynthetic control in low-light WT cultures, we noticed that WT cells responded to some extent differently after a 700-ms dark period when they were grown in moderate light, i.e. 40 μmol photons/m^2^/s. Therefore, the moderate light cultures from Figs. 1 and 2 were also monitored after 700-ms darkness (Supplementary Fig. 2). As expected^20^, there was no net oxidation of *b*-haems in relation to the pre-flash level in oxic WT cells (Supplementary Fig. 2a). The transient electron transfer re-acceleration before 500-ms of illumination was missing after the short dark period (Supplementary Fig. 2b; cf. Fig. 1c). The major difference between anoxic cultures from the two light growth conditions was a less pronounced net *b*-haems oxidation phase in WT from moderate light (Supplementary Fig. 2c; cf. Fig. 3e) that coincided with a significantly slower onset of electron transfer in the control samples (Supplementary Fig. 2d; cf. Fig. 3f). Apart from the missing AA effect, *stt7-1* controls resembled WT on the levels of *b*_6_*f* kinetics and electron transfer rates (Supplementary Figs. 2e and 2f). This stood in contrast to the above-mentioned *stt7-1* phenotype after 30-s darkness. Fig. 4d shows the *k*_*f*-red_ after 700-ms dark in moderate light cultures and, when compared with Fig. 2d (showing *k*_*f*-red_ after 30-s dark in those cultures), a strong photosynthetic control resulted in slowdown of *k*_*f*-red_ for all control samples. For instance, *k*_*f*-red_ in oxic WT was slowed down from ~350 s^−1^ to ~140 s^−1^, i.e. by a factor of 2.5 when cells dark-relaxed for 700-ms. Accordingly, the higher *k*_*f*-red_ in anoxic WT after 30-s resembled the invariably low anoxic *stt7-1* when the cells dark-relaxed for 700-ms only. This could explain the disappearing differences between WT and *stt7-1* on the electron transfer rates after 700-ms darkness (Supplementary Figs. 2d and 2f).

## Discussion

The *b*_6_*f* has been initially suggested as the AA-sensitive FQR component^26, 27, 29, 36^ which was later revised^24^. The important discovery of the PGR5 interactome^22, 37^ recognized PGR5/PGR5-Like1 as the AA-sensitive FQR for PGR5-dependent CEF, suggested by *in vitro* measurements. Here, we presented *in vivo* evidence that the cellular redox poise may equally dismiss and qualify the *b*_6_*f* as AA-sensitive FQR component in the same biological sample (Fig. 1). Although we also observed mild AA effects in the chloroplast that were associated with respiration inhibition in oxic samples (see also Supplementary Text 1), the contrasting AA effect on the *b*_6_*f* in anoxic conditions could explain the above-mentioned conflicting results when hunting for the AA-sensitive FQR in isolated chloroplasts. More specifically, the strongly reduced PQ pool activated the Stt7 kinase which itself was required for the AA-sensitive *b*_6_*f* feature (Fig. 2). Our data in Figs. 3 and 4 ruled out that PGR5 was responsible for both AA sensitivity of the b6f, and efficient electron transfer at the very onset of photosynthesis.

This challenges the above-mentioned PGR5/PGR5-Like1 model^23^. Instead, a recent proposal is substantiated^20^ which links the PGR5 polypeptide to the *b*-haems redox reactions in CEF-favouring conditions. It was suggested that PGR5 facilitates access to stromal electrons for the *b*_6_*f* Qi-site. In this scenario, supported by biochemical data^21^, PGR5 could be involved in efficient FNR recruitment to the thylakoid membrane (on the levels of PSI and *b6f*). Moreover, the *b*_6_*f* was proposed to operate in two types of Q cycles^20^: a canonical type during LEF and a Fd-assisted Q cycle during CEF. The latter Q cycle concept has been introduced elsewhere and early reports tied it to AA inhibition of Fd-dependent cyclic photophosphorylation^29, 36, 38, 39^. The dysfunctionality of the *pgr5 b*-haems redox reactions became more apparent in Figs. 3 and 4 when only a short dark period was given for equilibration before the flash-induced *b*_6_*f* turnover. Only WT showed a fast *b*-haems net oxidation (Fig. 3e) and we interpret the flat *pgr5 b*-haems signals (Fig. 3g) as an inefficient replenishment of reduced Fd to assist in the modified Q cycle. Accordingly, a pre-reduced haems *b*_h_^red^/*C*_i_^red^ couple in anoxic WT may produce two PQH_2_ at the Qi-site per Qo-site turnover after a flash by utilizing Fd^20^. Only a fraction of WT *b*_6_*f* might be engaged in this reaction after 700-ms darkness, considering the small amplitude compared to 30-s dark (cf. Figs. 3a and 3e). The 700-ms *b*_6_*f* kinetics in *pgr5* (Fig. 3g) are unlikely to be linked to enhanced photosynthetic control (lowering *k*_*f*-red_, Fig. 4c) but rather to inefficient *b*-haems oxidation in anoxic conditions. The latter eventually hampers the FeS domain movement of the Rieske ISP to prevent intrinsic short-circuits in the high-potential chain^40, 41^. Moreover, the diminished pmf in the mutant under these conditions^20^, evidenced by the lower steady state electron transfer rates (Fig. 3), was not in favour of enhanced photosynthetic control in *pgr5*. Another observation that argued against this was the WT-like ATP synthase activity upon a flash (Supplementary Fig. 2). Fd shortage in *pgr5* may be less severe after prolonged dark-relaxation and, on average, the *b*_6_*f* population produced only a small, yet significant *k*_*b*-ox_ difference after 30-s darkness (Fig. 4b). Under these conditions, the observed high initial electron transfer in anoxic *pgr5* resembled WT at least up to 10-ms of illumination (Figs. 3b and 3d). The comparable initial rates might be distantly related to findings in Arabidopsis *pgr5* showing WT-like CEF during the induction phase, despite a different CEF regulation capacity in *pgr5*^42^. It was demonstrated in Chlamydomonas that Fd-binding proteins PETO and ANR1 interact with the b6f, and at least PETO has access to the stromal *b*_6_*f* region^12, 43, 44^. If these interactions are PGR5-independent, a pool of Fd may be retained in proximity of the algal Qi-site by these auxiliary CEF proteins. Thus, despite modified FNR recruitment in *pgr5*^21^, the auxiliary Fd-binding proteins could accommodate CEF for a limited number of turnovers which resulted in WT-like initial electron transfer rates after a longer dark equilibration period.

It is important to note that our study only indirectly addressed CEF efficiency, judged from the AA inhibition of total electron transfer. We did not discriminate between LEF and CEF with intent (see also below). Although AA binding/unbinding may occur during continuous illumination, as suggested for other *b*_6_*f* Qi-site inhibitors^45^, electron transfer transients in the presence of AA should be mainly a function of LEF efficiency. The transients depended on the number of electrons accumulated in the chain during the 700-ms/30-s dark interval and the number of the active (AA-insensitive) *b*_6_*f* in the sample. A weakness of CEF estimations by using AA is that LEF efficiency may vary in controls and AA samples on the level of PSII. In theory, PSII acceptor side limitation is modulated upon disproportionation of PQ reduction by PSII and PQH_2_ oxidation by the diminished functional *b*_6_*f* population (in the *presence* of AA). The functional *b*_6_*f* population (in the *absence* of AA) is lower in *pgr5* under CEF-favouring conditions so that this disproportionation may be an intrinsic mutant feature, resulting in low LEF efficiency (Fig. 3) and a higher proportion of closed PSII centres^20^.

Here, we report for the first time that the *b*_6_*f* represented a bottleneck in anoxic *stt7-1* as well, which was possibly linked to a modified AA affinity in the Qi-site. When comparing with WT, the lower *b*_6_*f* activity in *stt7-1* could result from an inefficient Qi-site turnover (in analogy to *pgr5* above). Since the *stt7-1* steady state electron transfer rate was lower than in WT (independently of the dark adaptation period), the prevailing ΔpH was presumably higher in WT after 700-ms darkness. Considering the significant photosynthetic control in anoxic WT (Figs. 2d and 4d), the 700-ms behaviour of WT was disregarded to interpret *stt7-1* since the majority of the ΔpH was expectedly consumed only after 30-s darkness. Accordingly, the diminished electron transfer at the very onset of light was more pronounced when *stt7-1* cells dark-relaxed for 30-s (cf. Figs. 1f and 2c) instead of 700-ms (cf. Supplementary Figs. 2d and 2f). In contrast to *pgr5* with insufficient access to Fd at the Qi-site, we propose that the effect in anoxic *stt7-1* was due to incomplete switching to the Fd-assisted Q cycle. Stt7 interacts with the b6f^9^ and phosphorylates the complex directly^10^. Compared to *pgr5*, absence of the kinase may have more severe consequences on the *b*_6_*f* function in the dark-relaxed state. Therefore, a significant slowdown of *k*_*f*-red_ in addition to *k*_*b*-ox_ was observed in anoxic conditions after 30-s darkness (cf. Figs. 2d and 4a), as well as a significant drop of electron transfer in the first 10-ms of light (cf. Figs. 2c and 3d). This distinguished *stt7-1* from *pgr5* in addition to the AA effect. Since *stt7-1* was not fully AA resistant, it could be that a certain AA susceptibility was imposed by our conditions or that additional reactions were contributing to the *b*_6_*f* modulation, which could also be a function of the PQ pool redox state.

Nevertheless, we link these Stt7-dependent *b*_6_*f* adjustments to more efficient (total) electron flow which is AA-sensitive and the major contributing process during the first 100-ms of light (cf. Figs. 1f and 2c). This initial phase is presumably dominated by CEF although a recent analysis in the Chlamydomonas *stt7-9* mutant revealed that CEF was not affected in the mutant^15^. It should be noted that *stt7-9* is a leaky strain, unlike *stt7-1* used here^10^. Thus, *stt7-9* still contained the phosphorylation(s) in the *b*_6_*f* that might be responsible for the AA sensitivity^10^. We showed that candidate phosphorylation sites are found in the stromal region of the interacting subunit-IV and PETO, the loosely attached CEF effector^10, 11, 12, 16^. Furthermore, the CEF rates in *stt7-9* were obtained in the presence of DCMU^15^. The major methodical bottleneck of CEF measurements is that CEF shares several electron carriers with LEF. One disentanglement strategy is to abolish CEF by inhibitors such as AA. Potential drawbacks on LEF efficiency have been outlined above. Another strategy to measure CEF is to abolish LEF, using PSII inhibitors such as DCMU. In our study, it was intended to benefit from the PQ reductase activity of PSII to ensure maximal PQ pool reduction throughout the illumination regime in anoxic cells. Accordingly, we did not make further attempts to disentangle LEF and CEF since we also noticed in our conditions that PSII activity contributed to AA sensitivity of the *b*_6_*f* (*b*-phase in Fig. 1e) and suppression of electron flow in the presence of AA (Fig. 1f). This may reflect the PSII-dependent redox poise during CEF which has been recognized and discussed elsewhere^46, 47^. Therefore, AA-sensitive CEF measured in the absence of PSII photochemistry (or in PSI-specific far-red light) bears the risk of underestimating true maximal (PSII-poised) rates. In the absence of PSII activity, an increased AA-insensitive *b*_6_*f* population will eventually feed on the FQR activity of the proportionally diminished AA-sensitive *b*_6_*f* population, which should be in favour of non-cyclic processes to compete for the electron downstream of PSI.

Our data suggests that AA ligation depends on Stt7-mediated Qi-site adjustments. It remains to be investigated whether AA ligation resembles the situation in the respiratory cytochrome bc1 complex. This ambitious, yet auspicious task might shed new light on the haem *c*_i_ which, according to the structures^5, 6^, blocks the access for AA to ligate to haem *b*_h_ in a bc1-like manner^25^. One could envision that Stt7-dependent phosphorylation rearranges the Qi-site microenvironment, involving at least subunit-IV and/or PETO. The apparent Stt7-dependent AA effect, which we assume to rely on AA ligation to haem *b*_h_, might also involve slight coordination adjustments of haem *c*_i_ that facilitate stromal access to electrons and pave the way for bc1-like AA binding in the Qi-site. Stromal access for haem *c*_i_ is blocked in the structure by Arg95 in PetN and Asp35 in subunit-IV^5^, which are in proximity to the phosphorylated Thr4 in subunit-IV^10^ and the interface with the CEF effector PETO^12^.

To summarize, our tentative model proposes that the PGR5-dependent FNR recruitment regulation ensures the continuous supply of Fd for the *b*_6_*f* under conditions where AA-sensitive electron transfer dominates. Stt7-dependent *b*_6_*f* structure/function modifications are required to utilize these electrons for efficient Qi-site turnover. How these electrons may enter the complex, whether this step involves CEF supercomplexes^13, 14, 15, 48^ and which additional proteins are involved needs to be further characterized.

## Methods

### Cell Growth Conditions

Chlamydomonas reinhardtii WT t222+, stt7-1^49^, and pgr5^33^ were maintained at 20 μmol photons/m^2^/s on agar-supplemented Tris-acetate-phosphate plates^50^. When growing cells for experiments, liquid Tris-phosphate medium was devoid of acetate (TP). WT and stt7-1 cultures were grown at 40 μmol photons/m^2^/s (16 h light/8 h dark) and were bubbled with sterile air at 25 °C while shaking at 120 rpm (Figs. 1 and 2). The growth light for WT and pgr5 was set to 10 μmol photons/m^2^/s at otherwise identical conditions (Fig. 3). Grown cultures were diluted ~6-fold at least once after inoculation and grown to a density of ~2 × 10^5^ cells/ml before harvesting (5000 rpm, 5 min, 25°C). Cells were resuspended at 20 μg chlorophyll/ml in TP supplemented with 20% (w/v) Ficoll. Oxic samples were resuspended throughout the measurements in 2-ml open cuvettes. For oxygen-deprived conditions, cells were supplemented with 50 mM glucose, 10 U glucose oxidase and 30 U catalase in the cuvette, and then overlaid with mineral oil for 20 min in the dark. The enzyme stock solutions to obtain anoxic samples were freshly prepared.

### *In Vivo* Spectroscopy

Both oxic and anoxic samples were light-adapted in the cuvette for 20 min before the measurements of the electrochromic shift (ECS) and the cytochrome *b*_6_*f* redox kinetics. The measurements were obtained using a JTS-10 (Biologic) and are described in detail elsewhere^20^. In brief, all absorption changes were normalized to the ECS ΔI/I signals that were produced (<300 μs) after a saturating laser flash in the presence of 1 mM hydroxylamine (HA, from 1 M aqueous stock) and 10 μM 3-(3,4-dichlorophenyl)-1,1-dimethylurea (DCMU, from 10 mM ethanolic stock). Thus, HA and DCMU were used as PSII inhibitors to obtain the density of active PSI centres in the sample (measured as 1 charge separation/PSI). For *b*_6_*f* measurements, the flash intensity was lowered (to avoid double turnovers) which can be seen in Fig. 1b. There, both photosystems (with a ratio of ~1:1) were actively trapping photons and, when referring to the light-induced ECS amplitude (first records at 0.3-ms), the flash produced an accumulated ~0.7 charge separations/PSI. Once one photon trap (PSII) was inhibited in the presence of AA, about half the PSI population was excited by the flash (grey symbols in Fig. 1b). LEDs emitting ~150 μmol photons/m^2^/s of 630-nm actinic light (AL) were used for light adaptation in the cuvette. The AL was interrupted for 34-s during which two sub-saturating laser flashes were fired at 700-ms and 30-s. After these flashes, cytochrome *f* (554-nm – 0.27×(563-nm – 546-nm)) and the haems *b*_l_/*b*_h_ (563-nm) were measured, using a baseline drawn between 573-nm and 546-nm^20^. The interference filters were altered in a fixed order and the first set of records was dismissed after continuous AL adaptation to account for the altered light/dark regime. AL was on for ~1-min in between each dark period-containing measurement sequence. For calculations, means of six technical replicates were used for each wavelength. The dark phases were fit with a single-exponential function after the flash to obtain rate constants for cytochrome *f* reduction (*k*_*f*-red_, fit from 0.3-ms to 100-ms in oxic samples and 1-ms to 100-ms in anoxic samples) and *b*-haems oxidation (*k*_*b*-ox_, from 1-ms to 500-ms which had to be shortened in some anoxic samples owing to an established dark re-reduction phase). The standard errors of *k*_*f*-red_ and *k*_*b*-ox_ were comparable for each biological replicate whereas inhibition reduced the fit quality occasionally (Supplementary Fig. 3). Antimycin-A (AA, Sigma, Lot#061M4063V from 2011 and Lot#079M4102V from 2020 were used) was prepared freshly as 40 mM ethanolic stock and incubation at 40 μM in AL was for 30-min. Where indicated, these cells were illuminated for another 20-min in the absence of PSII photochemistry. The ECS (520-nm – 546-nm) was also deconvoluted in the dark pulse method where AL was shuttered briefly during a 10-s AL induction phase^39, 51, 52^. As for the *b*_6_*f* dark periods above, AL-adapted cells were measured after 700-ms and 30-s darkness (four technical replicates with 1-min AL in between). Two separate measuring sequences were used to avoid accumulating effects of shortly spaced dark intervals at the beginning of the induction phase^51^. AL was switched off at *t* = 0-ms in the sequences so that the linear ECS slopes before (S_L_ from – 4-ms to −1-ms) and after darkness (S_D_ from +1-ms to +5-ms) gave electron transfer rates before darkness (S_L_ – S_D_, expressed in separations/PSI/s). Four detecting pulses, spaced by 1-ms, recorded the ECS for slope calculations. The shown rates at 1-ms illumination were calculated from separate slopes obtained during 2-ms AL in light-adapted cells after 30-s dark (15 replicates spaced by 5-s).

## Acknowledgments

M.H. acknowledges support from Deutsche Forschungsgemeinschaft (DFG) Grant HI 739/13-1 and Grant HI 739/13-2. M.H. also acknowledges support from the RECTOR program (University of Okayama, Japan).

## References

1. Cramer WA, Hasan SS. Structure-function of the cytochrome *b*_6_*f* lipoprotein complex. In: Cytochrome complexes: Evolution, structures, energy transduction, and signaling (ed^(eds Cramer WA, Kallas T). Springer Netherlands (2016).

2. Saif Hasan S, Yamashita E, Cramer WA. Transmembrane signaling and assembly of the cytochrome b6f-lipidic charge transfer complex. Biochim Biophys Acta 1827, 1295–1308 (2013).

3. Alric J, Pierre Y, Picot D, Lavergne J, Rappaport F. Spectral and redox characterization of the heme ci of the cytochrome *b*_6_*f* complex. Proc Natl Acad Sci U S A 102, 15860–15865 (2005).

4. Lavergne J. Membrane potential-dependent reduction of cytochrome *b*_6_ in an algal mutant lacking photosystem I centers. Biochim Biophys Acta 725, 25–33 (1983).

5. Stroebel D, Choquet Y, Popot JL, Picot D. An atypical haem in the cytochrome *b*_6_*f* complex. Nature 426, 413–418 (2003).

6. Kurisu G, Zhang H, Smith JL, Cramer WA. Structure of the cytochrome *b*_6_*f* complex of oxygenic photosynthesis: tuning the cavity. Science 302, 1009–1014 (2003).

7. Rochaix JD. Redox regulation of thylakoid protein kinases and photosynthetic gene expression. Antioxid Redox Signal 18, 2184–2201 (2013).

8. Zito F, Finazzi G, Delosme R, Nitschke W, Picot D, Wollman FA. The Qo site of cytochrome *b*_6_*f* complexes controls the activation of the LHCII kinase. EMBO J 18, 2961–2969 (1999).

9. Dumas L, et al. A stromal region of cytochrome *b*_6_*f* subunit IV is involved in the activation of the Stt7 kinase in *Chlamydomonas*. Proc Natl Acad Sci U S A 114, 12063–12068 (2017).

10. Bergner SV, et al. STATE TRANSITION7-dependent phosphorylation is modulated by changing environmental conditions, and its absence triggers remodeling of photosynthetic protein complexes. Plant Physiol 168, 615–634 (2015).

11. Lemaire C, Girard-Bascou J, Wollman F-A, Bennoun P. Studies on the cytochrome *bdf* complex. I. Characterization of the complex subunits in *Chlamydomonas reinhardtii*. Biochim Biophys Acta 851, 229–238 (1986).

12. Buchert F, Hamon M, Gabelein P, Scholz M, Hippler M, Wollman FA. The labile interactions of cyclic electron flow effector proteins. J Biol Chem 293, 17559–17573 (2018).

13. Iwai M’ Takizawa K, Tokutsu R, Okamuro A, Takahashi Y, Minagawa J. Isolation of the elusive supercomplex that drives cyclic electron flow in photosynthesis. Nature 464, 1210–1213 (2010).

14. Terashima M, et al. Calcium-dependent regulation of cyclic photosynthetic electron transfer by a CAS, ANR1, and PGRL1 complex. Proc Natl Acad sci U S A 109, 17717–17722 (2012).

15. Takahashi H, Clowez S, Wollman FA, Vallon O, Rappaport F. Cyclic electron flow is redox-controlled but independent of state transition. Nat Commun 4, 1954 (2013).

16. Takahashi H, et al. PETO Interacts with Other Effectors of Cyclic Electron Flow in Chlamydomonas. Mol Plant 9, 558–568 (2016).

17. Clark RD, Hawkesford MJ, Coughlan SJ, Bennett J, Hind G. Association of ferredoxin-NADP^+^ oxidoreductase with the chloroplast cytochrome *b-f* complex. FEBS Lett 174, 137–142 (1984).

18. Zhang H, Whitelegge JP, Cramer WA. Ferredoxin:NADP^+^ oxidoreductase is a subunit of the chloroplast cytochrome *b*_6_*f* complex. J Biol Chem 276, 38159–38165 (2001).

19. Okutani S, et al. Three maize leaf ferredoxin:NADPH oxidoreductases vary in subchloroplast location, expression, and interaction with ferredoxin. Plant Physiol 139, 1451–1459 (2005).

20. Buchert F, Mosebach L, Gäbelein P, Hippler M. PGR5 is required for efficient Q cycle in the cytochrome *b*_6_*f* complex during cyclic electron flow. Biochem J 477, 1631–1650 (2020).

21. Mosebach L, et al. Association of Ferredoxin:NADP(+) oxidoreductase with the photosynthetic apparatus modulates electron transfer in Chlamydomonas reinhardtii. Photosynth Res 134, 291–306 (2017).

22. Munekage Y, Hojo M, Meurer J, Endo T, Tasaka M, Shikanai T. PGR5 is involved in cyclic electron flow around photosystem I and is essential for photoprotection in Arabidopsis. Cell 110, 361–371 (2002).

23. Hertle AP, et al. PGRL1 is the elusive ferredoxin-plastoquinone reductase in photosynthetic cyclic electron flow. Mol Cell 49, 511–523 (2013).

24. Moss DA, Bendall DS. Cyclic electron transport in chloroplasts. The Q-cycle and the site of action of antimycin. Biochim Biophys Acta 767, 389–395 (1984).

25. Huang LS, Cobessi D, Tung EY, Berry EA. Binding of the respiratory chain inhibitor antimycin to the mitochondrial *bc*_1_ complex: a new crystal structure reveals an altered intramolecular hydrogen-bonding pattern. J Mol Biol 351, 573–597 (2005).

26. Slovacek RE, Crowther D, Hind G. Cytochrome function in the cyclic electron transport pathway of chloroplasts. Biochim Biophys Acta 547, 138–148 (1979).

27. Tagawa K, Tsujimoto HY, Arnon DI. Role of chloroplast ferredoxin in the energy conversion process of photosynthesis. Proc Natl Acad Sci U S A 49, 567–572 (1963).

28. Tagawa K, Tsujimoto HY, Arnon DI. Separation by Monochromatic Light of Photosynthetic Phosphorylation from Oxygen Evolution. Proc Natl Acad Sci U S A 50, 544–549 (1963).

29. Arnon DI, Tsujimoto HY, McSwain BD. Ferredoxin and photosynthetic phosphorylation. Nature 214, 562–566 (1967).

30. Bailleul B, et al. Energetic coupling between plastids and mitochondria drives CO2 assimilation in diatoms. Nature 524, 366–369 (2015).

31. Joliot P, Joliot A. Quantification of the electrochemical proton gradient and activation of ATP synthase in leaves. Biochim Biophys Acta 1777, 676–683 (2008).

32. Velthuys BR. A third site of porton translocation in green plant photosynthetic electron transport. Proc Natl Acad Sci U S A 75, 6031–6034 (1978).

33. Johnson X, et al. Proton gradient regulation 5-mediated cyclic electron flow under ATP- or redox-limited conditions: a study of ΔATPase *pgr5* and ΔrbcL *pgr5* mutants in the green alga *Chlamydomonas reinhardtii*. Plant Physiol 165, 438–452 (2014).

34. Jans F, et al. A type II NAD(P)H dehydrogenase mediates light-independent plastoquinone reduction in the chloroplast of *Chlamydomonas*. Proc Natl Acad Sci U S A 105, 20546–20551 (2008).

35. Houille-Vernes L, Rappaport F, Wollman FA, Alric J, Johnson X. Plastid terminal oxidase 2 (PTOX2) is the major oxidase involved in chlororespiration in Chlamydomonas. Proc Natl Acad Sci U S A 108, 20820–20825 (2011).

36. Furbacher PN, Girvin ME, Cramer WA. On the question of interheme electron transfer in the chloroplast cytochrome *b*_6_ *in situ*. Biochemistry 28, 8990–8998 (1989).

37. DalCorso G, et al. A complex containing PGRL1 and PGR5 is involved in the switch between linear and cyclic electron flow in Arabidopsis. Cell 132, 273–285 (2008).

38. Binder RG, Selman BR. Ferredoxin Catalyzed Cyclic Photophosphorylation - Reversal of Dibromothymoquinone Inhibition by N,N,N’,N’-Tetramethyl-Para-Phenylenediamine. Z Naturforsch C 33, 261–265 (1978).

39. Joliot P, Joliot A. Cyclic electron transfer in plant leaf. Proc Natl Acad Sci U S A 99, 10209–10214 (2002).

40. Mulkidjanian AY. Ubiquinol oxidation in the cytochrome *bc*_1_ complex: Reaction mechanism and prevention of short-circuiting. Biochim Biophys Acta 1709, 5–34 (2005).

41. Allen JF. Cytochrome *b*_6_*f*. structure for signalling and vectorial metabolism. Trends Plant Sci 9, 130–137 (2004).

42. Nandha B, Finazzi G, Joliot P, Hald S, Johnson GN. The role of PGR5 in the redox poising of photosynthetic electron transport. Biochim Biophys Acta 1767, 1252–1259 (2007).

43. Peden EA, et al. Identification of global ferredoxin interaction networks in *Chlamydomonas reinhardtii*. J Biol Chem 288, 35192–35209 (2013).

44. Yang W, et al. Critical role of *Chlamydomonas reinhardtii* ferredoxin-5 in maintaining membrane structure and dark metabolism. Proc Natl Acad Sci U S A 112, 14978–14983 (2015).

45. Zito F, Alric J. Heme *c*_i_ or *c*_n_ of the Cytochrome *b*_6_*f* Complex, A Short Retrospective. In: Cytochrome Complexes: Evolution, Structures, Energy Transduction, and Signaling (ed^(eds Cramer WA, Kallas T). Springer Netherlands (2016).

46. Hauska GA, McCarty RE, Racker E. The site of phosphorylation associated with Photosystem I. Biochim Biophys Acta 197, 206–218 (1970).

47. Bendall DS, Manasse RS. Cyclic photophosphorylation and electron transport. Biochim Biophys Acta 1229, 23–38 (1995).

48. Steinbeck J, et al. Structure of a PSI–LHCI–cyt *b*_6_*f* supercomplex in *Chlamydomonas reinhardtii* promoting cyclic electron flow under anaerobic conditions. Proc Natl Acad Sci U S A 115, 10517–10522 (2018).

49. Depege N, Bellafiore S, Rochaix JD. Role of chloroplast protein kinase Stt7 in LHCII phosphorylation and state transition in Chlamydomonas. Science 299, 1572–1575 (2003).

50. Sueoka N. Mitotic replication of deoxyribonucleic acid in *Chlamydomonas reinhardi*. Proc Natl Acad Sci U S A 46, 83–91 (1960).

51. Nawrocki WJ, Bailleul B, Cardol P, Rappaport F, Wollman FA, Joliot P. Maximal cyclic electron flow rate is independent of PGRL1 in Chlamydomonas. Biochim Biophys Acta Bioenerg 1860, 425–432 (2019).

52. Bailleul B, Cardol P, Breyton C, Finazzi G. Electrochromism: a useful probe to study algal photosynthesis. Photosynthesis Res 106, 179–189 (2010).

