## Supplemental Data 1 for "Cytochrome *b*_6_*f* complex inhibition by antimycin-A requires Stt7 kinase activation but not PGR5"

***Title***:

***Author Line***:

Felix Buchert[1]* and Michael Hippler[1,2]*

***Author Affiliation***:

[1]Institute of Plant Biology and Biotechnology, University of Münster, Schlossplatz 8, 48143 Münster, Germany

[2]Institute of Plant Science and Resources, Okayama University, Kurashiki, Japan

**Contents:**

**Supplementary Text 1**

**Supplementary Figs. 1 – 3**

**Supplementary References**

**Supplementary Text 1** The respiratory inhibitor antimycin-A changed the redox state of the chloroplast slightly, which was described for oxic samples in panels a–c of Fig. 1 in the main text. The relatively high initial rates of the PSII-inhibited, antimycin-A containing samples were not fully unexpected under *semi*-oxic conditions in Fig. 1c of the main text. Before addition of the PSII inhibitors, cells were not experiencing an anoxic treatment despite the presence of the respiratory inhibitor antimycin-A. Considering the main focus of the manuscript, we did not further check whether PTOX-mediated chlororespiration was altered by preventing water splitting, thus failing to antagonize NDA2-mediated plastoquinone pool reduction during the 30-s dark period[1]. It was also not further tested whether a smaller $\Delta$pH in the dark facilitated initial electron rates via the $b_6f$ when PSII was inhibited. It is certain that ceasing electron transfer diminished both the (re-)generation rate of reduced thioredoxin to keep ATP synthase in the $\gamma$-dithiol state, and the driving force for ATP synthesis. These effects are expected to lower the proton motive force in the dark which is established by proton pumping via algal ATP synthase. The modulation of the algal proton motive force in the dark was discussed recently[2].

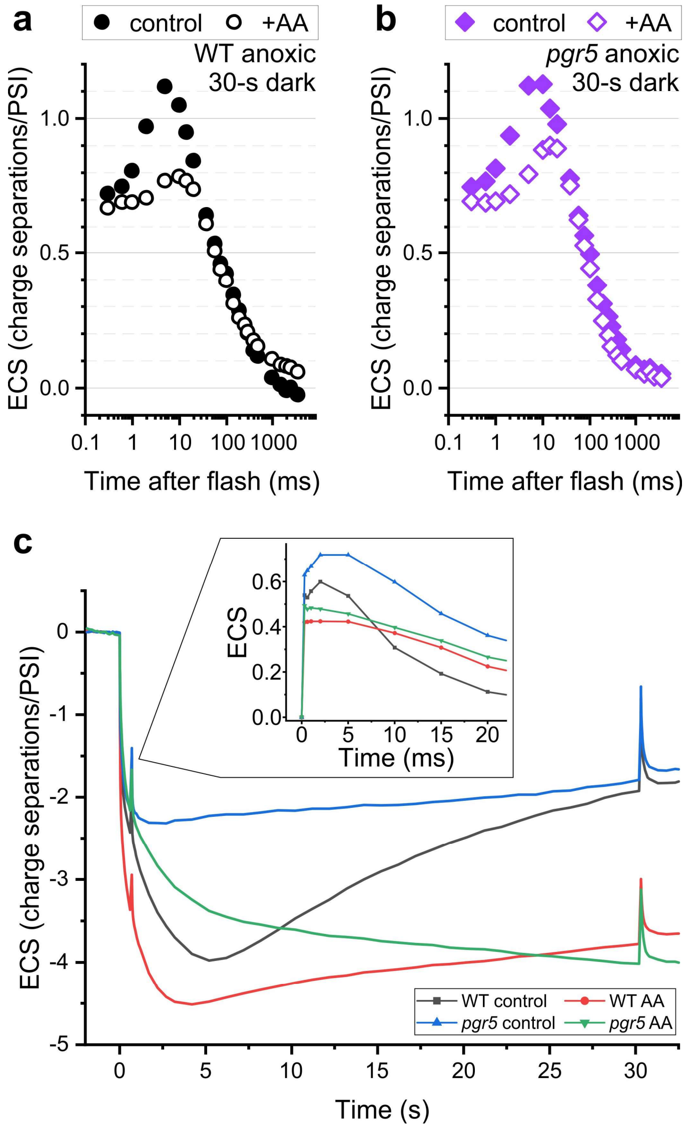


**Supplementary Fig. 1** The conditional sensitivity of the cytochrome $b_6f$ complex to antimycin-A is displayed in *pgr5* by measuring the electrochromic shift (ECS). The charge separation activity of the cytochrome $b_6f$ complex resulted in positive signals, developing in the 10-ms range after the flash in controls of **a** WT and **b** *pgr5*. Treatment with 40 µM antimycin-A (+AA, open symbols) resulted in a less pronounced activity during the 10-ms phase. **c** The full ECS traces during the measurement are shown and the developing signals during the first flash, after 700-ms darkness, are magnified (inset). The full traces reference was the ECS before darkness ($t < 0$) and the inset signals were referenced to the pre-flash ECS. The disappearing cytochrome $b_6f$ complex activity is seen in both strains by the absence of the 10-ms phase in the presence of AA (inset). The flash kinetics after 30-s darkness are shown in panels a and b (after rotating/correcting the curves for a linear dark drift between 27-s and 30-s).

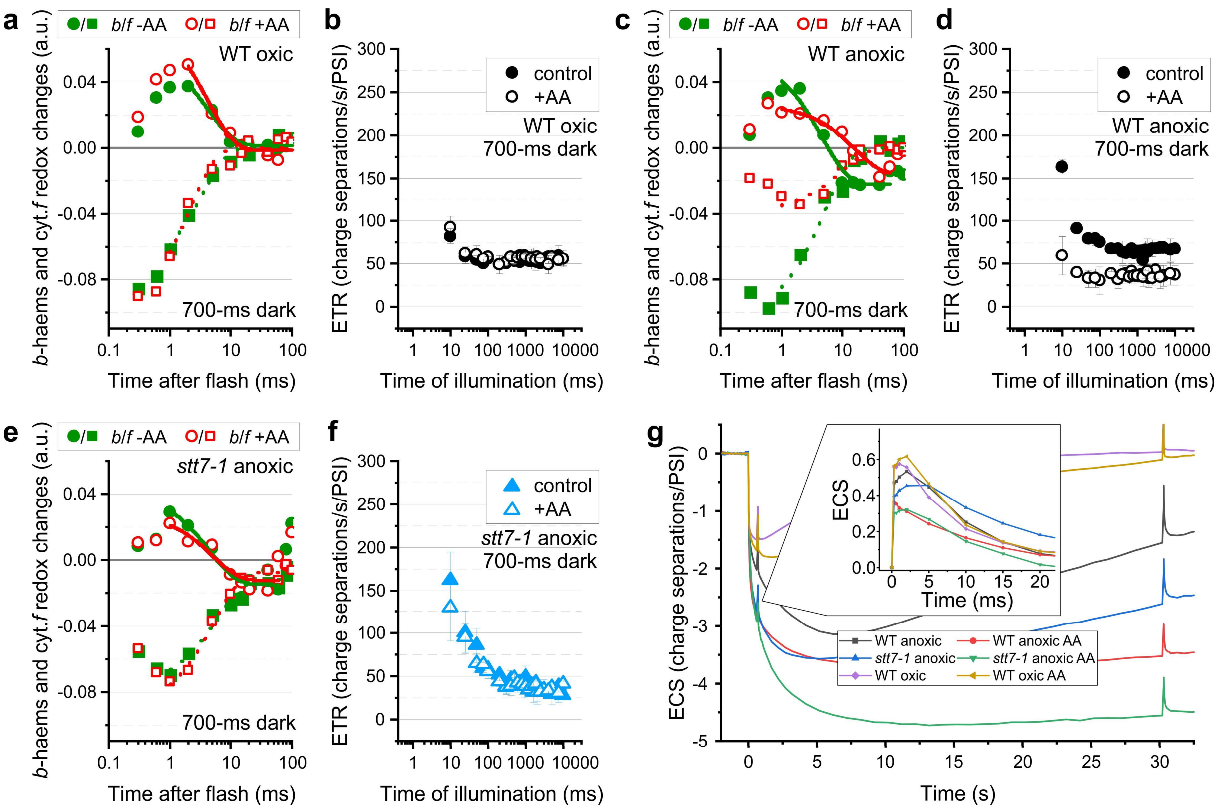


**Supplementary Fig. 2** The Stt7-dependent antimycin-A (AA) sensitivity of the cytochrome $b_6f$ complex and photosynthetic electron transfer are shown in samples that experienced 700-ms darkness. Representative kinetics and, where indicated, means of three biological replicates (± SD) are shown. Refer to Figs. 1 and 2 in the main text for measurements after 30-s dark. **a** Redox signals for cytochrome *f* (cyt.*f*, squares) and *b*-haems (circles) are shown in oxic WT controls (-AA, closed symbols) and after treatment with antimycin (+AA, open symbols). The AA treatment did not substantially alter cyt.*f* re-reduction upon a flash, shown by a signal increase in the positive direction that finished no later than 20-ms. AA did also not affect the *b*-haems reduction (positive signals) and re-oxidation to pre-flash levels, finishing at ~50-ms at pre-flash levels. **b** Electron transfer rate (ETR) measurements after 700-ms darkness differed only by lacking a re-acceleration phase before 500-ms of light. No AA effect was observed. **c** The cytochrome $b_6f$ redox kinetics in anoxic WT differed by showing the inhibitory AA effect and control samples displayed a *b*-haems oxidation phase below the pre-flash levels. **d** The ETR relied on AA-sensitive processes when anoxic WT was re-illuminated after 700-ms darkness. **e** Anoxic *stt7-1* controls resembled WT from panel *C* but no comparable AA effect on the cytochrome $b_6f$ redox kinetics was measurable after 700-ms darkness. **f** The AA effect was also less pronounced during ETR measurements in anoxic *stt7-1* where initial rates were as in WT of panel *D* but levelled off below the WT steady state. **g** The full ECS traces during the measurement are shown and the developing signals during the first flash, after 700-ms darkness, are magnified (inset). The full traces reference was the ECS before darkness ($t < 0$) and the inset signals were referenced to the pre-flash ECS. The disappearing cytochrome $b_6f$ complex activity upon AA treatment is seen only in anoxic WT by the absence of the 10-ms phase that produced positive signals (red curve, inset). The flash kinetics after 30-s darkness are shown in Figs. 1 and 2 in the main text (after rotating/correcting the curves for a linear dark drift between 27-s and 30-s).

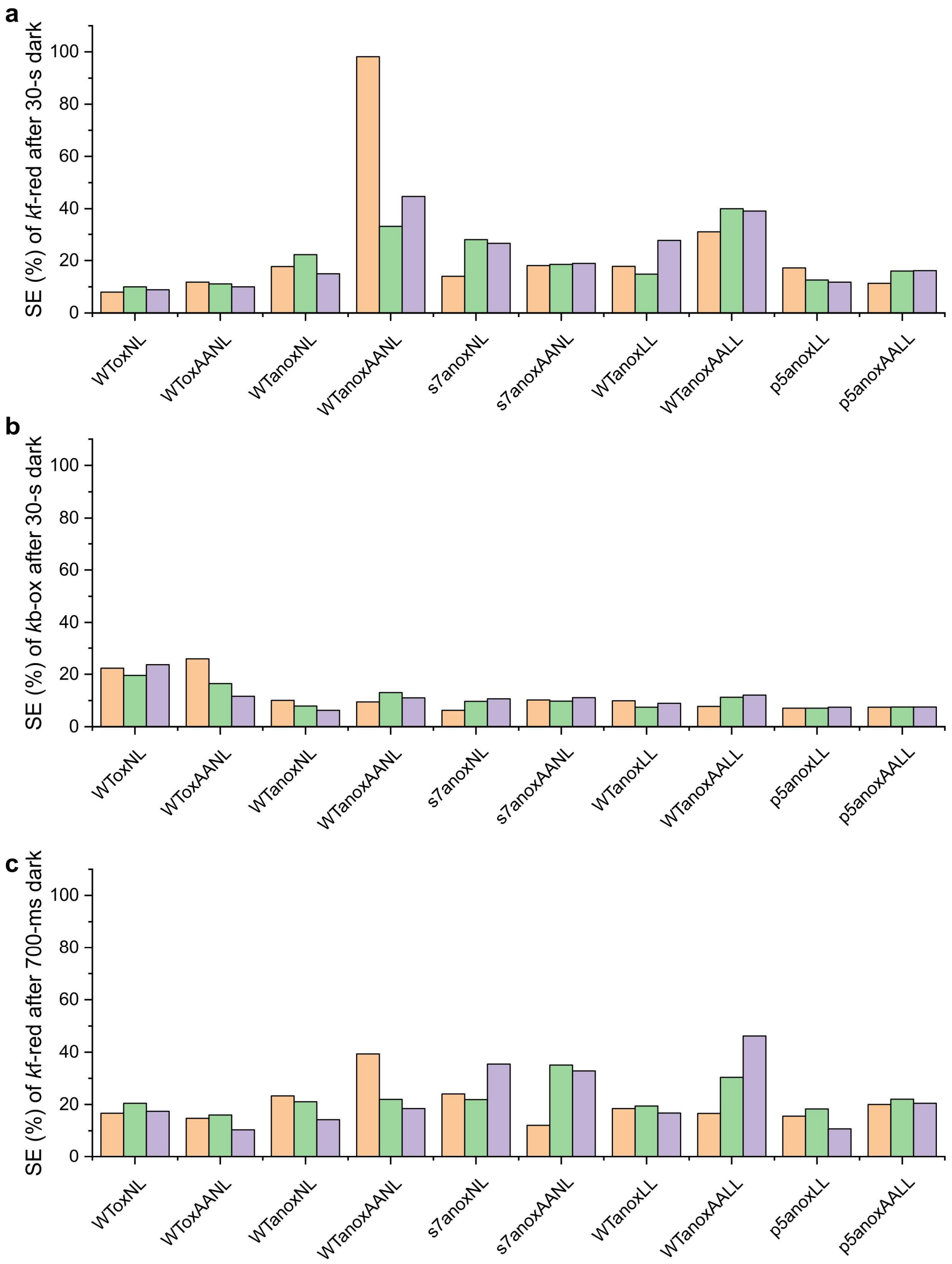


**Supplementary Fig. 3** The relative standard errors (SE, in %) are shown for **a** cytochrome *f* re-reduction (*k*f-red) and **b** *b*-haems oxidation (*k*b-ox) after 30-s darkness, as well as **c** *k*f-red after 700-ms darkness. See Materials and Methods in the main text which contains the means of these biological replicates in Figs. 2 and 4. Abbreviations are: WT, wild type; s7, *stt7-1*; p5, *pgr5*; ox, oxic; anox, anoxic; AA, antimycin-A; NL and LL, cultures grown at 40 and 10 μmol photons/m$^2$/s, respectively.

### Supplementary References


1. Nawrocki WJ, Tourasse NJ, Taly A, Rappaport F, Wollman FA. The plastid terminal oxidase: its elusive function points to multiple contributions to plastid physiology. *Annu Rev Plant Biol* **66**, 49-74 (2015).

2. Finazzi G, Drapier D, Rappaport F. Chapter 18 - The $CF_OF_1$ ATP Synthase Complex of Photosynthesis. In: *The Chlamydomonas Sourcebook (Second Edition)* (ed^(eds Harris EH, Stern DB, Witman GB). Academic Press (2009).
